# Genetic basis of lineage-specific evolution of fruit traits in hexaploid persimmon

**DOI:** 10.1101/2023.05.23.542005

**Authors:** Ayano Horiuchi, Kanae Masuda, Kenta Shirasawa, Noriyuki Onoue, Ryusuke Matsuzaki, Ryutaro Tao, Yasutaka Kubo, Koichiro Ushijima, Takashi Akagi

## Abstract

Frequent polyploidization events in plants have led to the establishment of many lineage-specific traits representing each species. Little is known about the genetic bases for these specific traits in polyploids, presumably due to plant genomic complexity and their difficulties of applying genetic approaches. Hexaploid Oriental persimmon (*Diospyros kaki*) has evolved specific fruit characters, including wide variations in fruit shapes and astringency. In this study, using whole-genome diploidized/quantitative genotypes from ddRAD-Seq data of 173 persimmon cultivars, we examined their population structures and potential correlations between their structural transitions and variations in nine fruit traits. The population structures of persimmon cultivars were highly randomized and not substantially correlated with the representative fruit traits focused on in this study, except for fruit astringency. With genome-wide association analytic tools considering polyploid alleles, we identified the loci associated with the nine fruit traits; we mainly focused on fruit-shape variations, which have been numerically characterized by principal component analysis of elliptic Fourier descriptors. The genomic regions that putatively underwent selective sweep exhibited no overlap with the loci associated with these persimmon-specific fruit traits. These insights will contribute to understanding of the genetic mechanisms by which fruit traits are independently established, possibly due to polyploidization events.

## Introduction

In contrast to animals, plant taxa have historically experienced frequent polyploidization events. Ancient paleogenome doubling (or paleo-polyploidization) often triggered establishment of novel traits in plants, in lineage-specific manners (Van de Peer et al. 2017), such as water adaptation in eelgrass (*Zostara marina*) (Olsen et al. 2016). More recent polyploidizations have also contributed to ecological adaptations, such as in the genus *Fragaria* (Liston et al. 2014; Wei et al. 2019), and crop domestication or improvement (Salman-Minkov et al. 2016; Akagi et al. 2022). Incremental yield increases (Heslop-Harrison and Schwarzacher 2007), conversion to self-pollinating systems (Hauck et al. 2005; Muñoz-Sanz et al. 2020; Masuda et al. 2022), and other traits thought to be beneficial for cultivation have been reported to involvement recent polyploidization (Akagi et al. 2022). Regarding fruit characters, ancient and recent polyploidization events have also played important roles in promoting the lineage-specific evolution. Fruit-ripening pathway in tomato (TGC 2012), acquisition of a specific pungent odorant in durian (Teh et al. 2017) and oil composition specific to olive fruits (Unver et al. 2017) were thought to be triggered by lineage-specific paleo-polyploidization events. Recent polyploidization events have often been involved in fruit development, classically represented by increases in fruit size (Langhe et al. 2009 for banana; Wu et al. 2012 for kiwifruit). However, their genetic mechanisms are not well understood due to genomic complexity and difficulties in polyploids (Akagi et al. 2022).

Cultivated Oriental persimmon (*Diospyros kaki*), a major fruit crop in East Asia, is autohexaploid with hexasomic inheritance (2n = 90) (Akagi et al. 2012). The genus *Diospyros* also includes other polyploid species, such as ebony (*D. ebenum*, hexaploid with 2n = 90) and American persimmon (*D. virginiana*, tetra- or hexaploid with 2n = 60, 90) (Tamura et al. 1998), which are mainly utilized as timber or edible fruits. The establishment of hexaploid *D. kaki* has given rise to many types of specific fruit traits, including variation in astringency (Akagi et al. 2011; Yamada and Sato 2016) and diverse fruit shapes (Maeda et al. 2018, 2019). Regarding astringency, the fruit of wild relatives close to *D. kaki* stably accumulates high levels of proanthocyanidins (or condensed tannins), which results in strongly astringent fruit. The loss of fruit astringency is specific to cultivated Oriental persimmon and is caused by two independent genetic factors (Sugiura and Tomana 1983; Akagi et al. 2011). One is dependent on volatile products (alcohol and acetaldehyde) emissions from the seeds, which insolubilizes tannin polymers for reducing astringency (defined as “volatile dependent group (VDG)” or “non-pollination-constant non-astringent (non-PCNA) type”). The other factor depends on reduction in the ability to accumulate proanthocyanidins, which is regulated by a single hypothetical locus named *ASTRINGENCY* (defined as “volatile independent group (VIG)” or “PCNA type”) (Akagi et al. 2011). In terms of fruit-shape variation, Oriental persimmon cultivars exhibit wide multidimensional diversity in their fruit shapes, which is not observed in close wild relatives (Maeda et al. 2018). Persimmon fruit-shape variations have been numerically characterized by principal component analysis (fruit shape PCA) using elliptic Fourier descriptors in SHAPE program (Iwata and Ukai 2002; Maeda et al. 2018). The SHAPE program was applied not only for evaluating fruit shapes but also for comparative evolutionary studies on leaf development among grape cultivars (Chitwood et al. 2016). Potentially, this program is useful in elucidating the unclear evolutionary process for fruit shape variations in hexaploid persimmon combined with genetic information. The physiological mechanism that determines fruit-shape variation involves persimmon-specific regulatory pathways using fruit shape PCA and transcriptome analysis, which are independent of genes involved in determining tomato fruit shape (Maeda et al. 2019). However, the genetic basis for the acquisition of fruit-shape variations in hexaploid persimmon is unclear. These Oriental-persimmon-specific traits are distributed widely in the current populations, while little is known about the transitions of population structures, or the breeding processes linked to these specific fruit traits. Insights into these processes would contribute to understanding of the genetic and evolutionary mechanisms, possibly associated with polyploidization events, that have led to the establishment of lineage-specific fruit traits.

Assessing the genetic architecture of traits with multiple alleles (and often mixed inheritance patterns) is challenging in polyploid species (Dufresne et al. 2013). Inheritance modes or chromosomal behaviors in polyploids are distinct from those in diploids. Genotyping in polyploids is complicated due to the possibility of there being more than two alleles at each locus and the existence of multiple heterozygous states. In the autohexaploidy inheritance mode of Oriental persimmon (Akagi et al. 2012; Masuda et al. 2020), we can define five heterozygous states: AAAAAa (pentaplex), AAAAaa (quadruplex), AAAaaa (triplex), AAaaaa (duplex), and Aaaaaa (simplex). Recent progress in large-scale sequencing technologies and development of analytical tools for polyploids, such as updog (for quantitative genotyping in polyploids; Gerard et al. 2018), StAMPP (for calculation of genetic distances with quantitative allelic states; Pembleton et al. 2013), and GWASpoly (for genome-wide association studies (GWAS) considering allele dosages; Rosyara et al. 2016), has allowed the study of genetics and genomics to interpret the evolution of variations in non-model polyploid crops. Using these advanced techniques, we attempted to study the genetic basis and evolutionary processes involved in the formation of Oriental-persimmon-specific fruit traits involved in recent polyploidization events, mainly from the perspective of population genetics in complicated hexaploid genomes.

## Materials and Methods

### Plant materials and phenotypes examined

Genomic DNA was extracted from young leaves of 173 persimmon cultivars sampled at the experimental orchards of Kyoto University, Kyoto, Japan and the Grape and Persimmon Research Station, National Institute for Fruit and Tea Science (NIFTS), Hiroshima, Japan (Supplementary Table S1) using the CTAB method. For phenotypes, we targeted nine fruit traits: four fruit-shape indexes (two major principal components in Maeda et al. 2018 and two fruit groove indexes, as described later), PCNA- and non-PCNA-type astringency loss (Akagi et al. 2011), fruit size, fruit-ripening time, and peel color, for which hexaploid persimmon cultivars exhibit notable diversity. For fruit shapes, we applied two principal components (fruit shape PC1 and PC4) with elliptical Fourier descriptors (EFDs) in the SHAPE program (Iwata and Ukai 2002) to fruit longitudinal sections (Fig. 1). which had previously been characterized as the main variations among the cultivars (Maeda et al. 2018). The fruit shape PC1 scores were strongly correlated with the ratio of length to diameter and fruit shape PC4 was associated with fruit flatness (Maeda et al. 2018). We also used the presence of grooves and the measured groove depths in the fruit peripheral regions. Fruit size, ripening time, peel color, and the two astringency types were annotated by quantitative categories according to a previous report (Fruit Tree Experiment Station of Hiroshima Prefecture, 1979) and the NIAS Genebank database (https://www.gene.affrc.go.jp/databases-plant_search_char.php). The fruit astringency in hexaploid persimmon was classified into four types based on their astringency-loss ability, depending on the presence of seeds or astringency: pollination-constant astringency (PCA), pollination-variant astringency (PVA), pollination-variant non-astringency (PVNA), and pollination-constant non-astringency (PCNA) (Sugiura and Tomana 1983). PCA and PCNA-type fruits are always astringency and non-astringency, respectively, regardless of the presence of seeds. For PCNA type astringency loss, four astringency types were classified into two categories, including PCNA and non-PCNA (PCA, PVA, and PVNA). For non-PCNA type astringency loss, quantitative values of astringency-loss ability were modified to 0, 1, 2 with PCA, PVA, PVNA, respectively (Akagi et al. 2011).

**Fig. 1.**
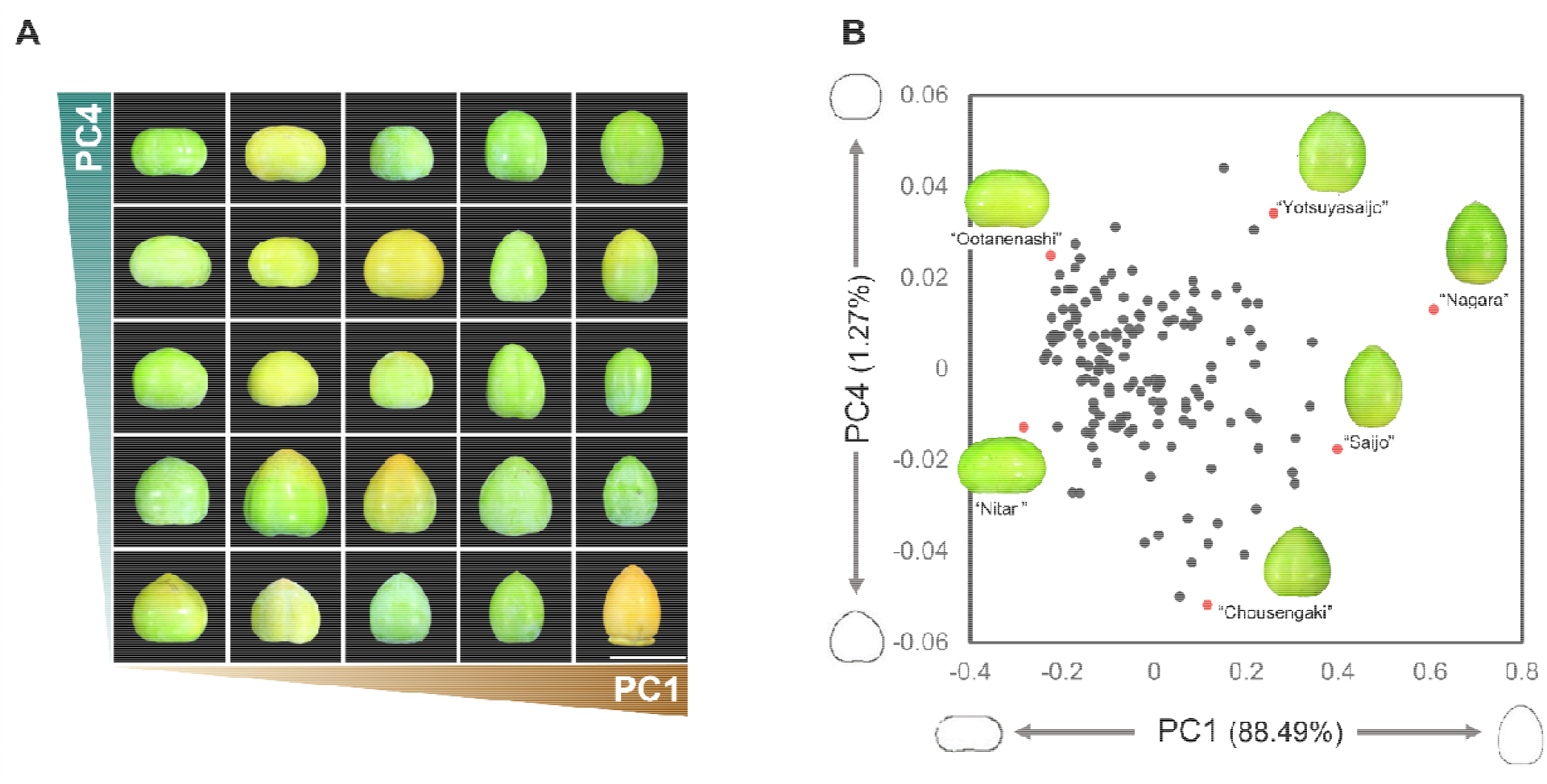
Fruit-shape diversity in persimmon cultivars. **(A)** Persimmon fruit-shape variations in the longitudinal direction, with various fruit-shape PC1/PC4 indexes. The length of the scale bar is 10 cm. **(B)** PC1 and PC4 in the previous principal component analysis for 153 persimmon cultivars (modified from Maeda et al. 2018). The percentage of variance explained by each principal component is provided in parentheses on each axis.

### Quantitative genotyping

ddRAD-Seq libraries were prepared with ∼200 ng gDNA, as described previously (Masuda et al. 2020). The libraries were sequenced on an Illumina HiSeq4000 platform, and PE100 reads were generated at the Vincent J. Coates Genomics Sequencing Laboratory, University of California Berkeley. The data from 173 persimmon cultivars were mapped to closely-relative species *Diospyros lotu*s (Yonemori et al. 2008, Nishiyama et al. 2018) whole-chromosome pseudomolecule sequences (Akagi et al. 2020) as the reference using the Burrows–Wheeler Aligner (BWA) (Li et al. 2009) with default settings. The resultant sam files were converted into vcf files with Samtools (Li and Durbin 2009) and VarScan v2.3.7 (Koboldt et al. 2012). The vcf files were filtered with the following thresholds for GWAS analysis (SNP nos=113,181): minor allele frequency (>0.01), min DP (>50), and max missing (=0.9). The files were further filtered with LD values (r^2^<0.3) for population structure analyses (SNP nos=16,956) using PLINK (Purcell 2007). The quantitative allele composition (0–6 or nulliplex–hexaplex) was determined by using updog, a genotyping program for polyploids (Gerard et al. 2018) and filtered by post-probability of genotyping (propmis<0.1). Note that the quantitative genotypes of a few nonaploid cultivars were determined as hexaploid cultivars.

### Population structure analysis

For ADMIXTURE analysis and evolutionary tree construction, we called diploidized genotypes with a custom Python-based program to set the hetero-tolerance to 5% (or 0/1 for 5%–95% alternative alleles), as described previously (Masuda et al. 2020). We converted diploidized vcf files to bed files with PLINK version 1.07 (Purcell 2012) for use with ADMIXTURE ver. 1.3.0 (Alexander et al. 2009). The *K* (number of ancestral populations) value was initially set from 2 to 10 to examine the most likely cluster numbers based on delta-K (ΔK) analysis (Evanno et al. 2005). Evolutionary topology was calculated by the NJ method using MEGA X (Kumar et al. 2018; 100 bootstrap replicates per branch); concatenated SNPs data were converted from vcf to phylip formats according to the modified method of https://github.com/edgardomortiz/vcf2phylip. Principal component analysis (PCA) was conducted using 16,956 SNPs with an error tolerance rate <10% with pcaMethods in R (Stacklies et al. 2007). Pearson linear correlations were calculated between PC1-PC20 and the nine fruit traits. The distance matrix of the genetic diversity considering hexasomic allele state, was calculated with StAMPP (Pembleton et al. 2013).

### Polyploidy GWAS and expression-based GWAS including allele-dosage contribution

The quantitative genotype data (nulliplex–hexaplex) in 150 of 173 persimmon cultivars, of which confident/sufficient phenotypic data were found in the legacy databases, were applied to GWASpoly Version 2 (Rosyara et al. 2016). GWASpoly was performed based on a linear mixed model with population structures using the K model considering a random polygenic effect. GWASpoly can consider various allele dosages with two types of models (Fig. 2): (i) an additive model, in which the contribution is assumed to increase with allele dosage, and (ii) a dominant model, in which the contribution is considered binary. For a dominant model, we considered simplex–triplex allele states for the threshold (1-dom for simplex, 2-dom for duplex, and 3-dom for triplex) (Fig. 2). After filtering using updog, a total of 113,181 SNPs were used for GWAS analysis. The boundaries for significance were set by Bonferroni-corrected threshold of 0.1 (-log(*P*-value) > 6.26 - 6.29). Local LD values were calculated for loci surrounding the detected GWAS peaks to confine the genetic regions with potential links to the peaks.

**Fig. 2.**
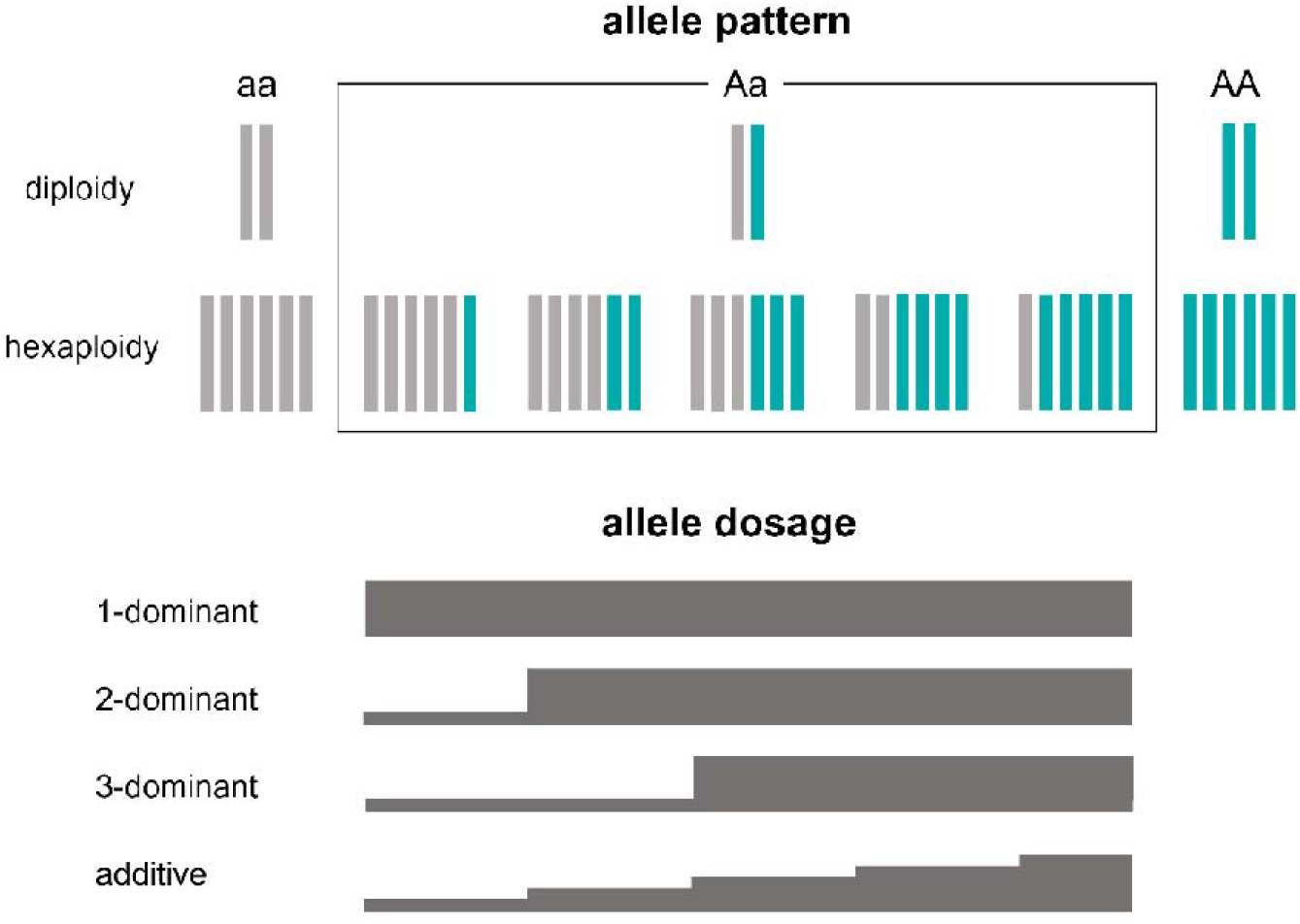
Definition of genotypes in hexaploidy allelic state. Five allelic states in hexaploid persimmon, AAAAAa (pentaplex), AAAAaa (quadruplex), AAAaaa (triplex), AAaaaa (duplex), and Aaaaaa (simplex), are uniformly defined as “heterozygous” in diploidized genotypes (A). GWASpoly can set thresholds of allele dosages in the dominant model and determine the quantitative effects of allele dosages in the additive model.

For expression-based GWAS (eGWAS), we targeted *KNOTTED1-LIKE FROM ARABIDOPSIS THALIANA* (*KNAT1*), *SEEDSTICK* (*STK*), and *cytokinin oxidase 5* (*CKX5*), which are thought to be important for persimmon fruit-shape diversity (Maeda et al. 2019). Reads per kilobase and million mapped reads (RPKM) values were applied as the response variables at the early fruit developmental stage critical for shape determination (late June, in Japan; Maeda et al. 2019). The mRNA-Seq data of 43 cultivars was obtained from Maeda et al. (2019). We newly sequenced additional mRNA-Seq libraries from another 22 persimmon cultivars using developing fruits at the same stage 4 (mid-June 2021) as in Maeda et al. (2019) (Supplementary Table S2), which are supposed to be important for the determination of fruit shape in persimmon cultivars (Maeda et al. 2018). Total RNA was isolated using the hot borate method, according to Maeda et al. (2019). Extracted total RNA was processed in preparation for Illumina Sequencing, as previously described (Masuda et al. 2020). In brief, mRNA was purified using the Dynabeads mRNA Purification Kit (Thermo Fisher Scientific, Waltham, MA, USA). The mRNA libraries were constructed using the KAPA RNA HyperPrep Kit (Roche, Basel, Switzerland). The libraries were sequenced on Illumina’s HiSeq 4000 (50-bp SR). All Illumina sequencing was conducted at the Vincent J. Coates Genomics Sequencing Laboratory at UC Berkeley. Raw sequencing reads were processed using Python scripts (https://github.com/Comai-Lab/allprep/blob/master/allprep-13.py) for preprocessing and demultiplexing of sequencing data. The mRNA-seq reads were aligned to the reference CDSs of *D. lotus* whole-genome sequences (Akagi et al. 2020) using the default parameters of the BWA (version 0.7.15) (Li and Durbin, 2009) (http://bio-bwa.sourceforge.net/). Normalized expression levels (RPKMs) of these important three genes were calculated and applied to GWASpoly in additive and dominant models.

### Test for site-frequency spectrum (SFS)-based selective sweeps

Selective sweep was estimated using a method based on the site-frequency spectrum (SFS) for 113,494 SNPs considering MAF and missing rates using 173 cultivars. Major methods to detect selective sweep would be categorized into SFS-based and extended haplotype-based (Nielsen et al. 2007). In polyploid species, detection of selective sweep would be limited with SFS-based, due to the difficulty in haplotype expectation with binary heterozygosity (*H*_*p*_), according to a previous report (Rubin et al. 2010). The window contains only the positions of genotypic data. The SFS-based method was applied in 100-kb steps with a 400-kb sliding window using pooled the three or fewer SNPs removed. The average expected heterozygosity was given for each window as follows:

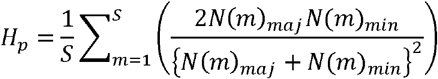

*S* is the number of SNP Positions in the window, and *N*(*m*)_maj_ and *N*(*m*)_min_ are the respective numbers of major and minor alleles at the *m*th SNP locus in the window. The individual *H*_*p*_values were then *Z*-transformed as follows: *ZH*_*p*_ = (*H*_*p*_ *− μH*_*p*_)/*σH*_*p*_, as previously reported for pooled heterozygosity in selective sweep analysis in chicken (Rubin et al. 2010).

## Results

### Population structure of persimmon cultivars

We attempted to uncover population structures in hexaploid persimmon using four methods considering quantitative allelic states: principal component analysis (PCA), ADMIXTURE, evolutionary topology, and genetic-distance matrixes with StAMPP (Pembleton et al. 2013). Both of PCA and ADMIXTURE analyses suggested weak clusterization coordinated with genotypes of astringency/non-astringency traits in fruit (Fig. 3 A–B, Supplementary Fig. S1). This result is consistent with the PCNA-type (or strictly J-PCNA; Akagi et al. 2011) astringency-loss group originating in a very narrow genetic background, such as ‘Gosho’ and related varieties (Yamada and Sato 2016), and the PCNA phenotype required recessive homozygosity at the single *ASTRINGENCY* locus (Akagi et al. 2011). However, in the PCA with genetic data, genome PC1 explained only 4.8% of the variance, suggesting random distribution during cultivar differentiation rather than subpopulations in peach (Akagi et al. 2016). Genetic-distance matrixes from StAMPP analysis, considering quantitative genotypes in hexasomic allele compositions, exhibited substantially low genetic correlations among the persimmon cultivars, which was consistent with the results of PCA and supported the random differentiation of persimmon cultivars (Fig. 3C).

**Fig. 3.**
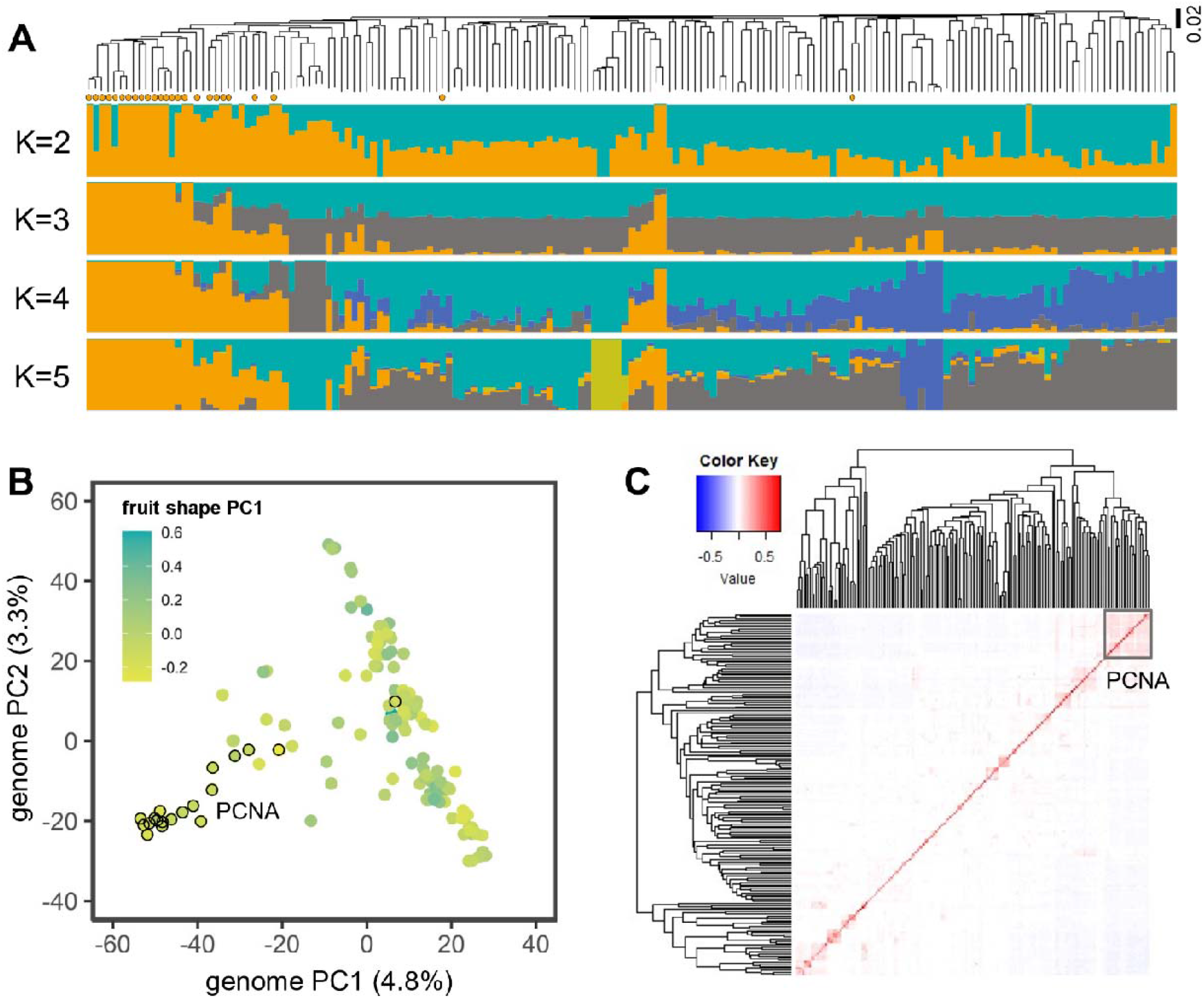
Population structures of persimmon cultivars. (A) Phylogenetic analysis and proportion of ancestry from K=2–5 inferred using ADMIXTURE in 173 hexaploid persimmon cultivars. The orange circles indicated PCNA cultivars. PCNA-type astringency-loss varieties mostly formed a separate cluster. The cultivar names were listed in Supplementary Table S1. (B) Distribution of the first two principal components (genome PC1 and PC2) in the PCA. Fruit-shape PC1 values (Maeda et al. 2018) are shown with a color gradient. No substantial correlation was detected between population structure and fruit-shape PC1. (C) Genetic distance matrix in 173 hexaploid persimmon cultivars created with StAMPP using quantitative genotypes.

Correlation tests between principal component (PC) values and fruit traits exhibited clear correlation between PCNA-type astringency loss and PC1 (*r*^2^=0.52; Fig. 4, Supplementary Fig. S2A), which was consistent with their co-segregation in the described structural analysis using ADMIXTURE or evolutionary topology (Figs. 3 and 4). However, there were no substantial correlations (*r*^2^<0.35) between the PC1–PC20 values and the other fruit phenotypes examined (Fig. 4). Only weak correlations (*r*^2^>0.1) were detected between non-PCNA-type astringency loss and the genome PC2 (*r*^2^=0.32), fruit shape PC1 and the genome PC7 (*r*^2^=0.10), presence of a groove and the genome PC10 (*r*^2^=0.12), and fruit size and the genome PC1 (*r*^2^=0.12) (Supplementary Fig. S2B-E). These results suggest that, except for PCNA-type astringency loss, notably diversified fruit traits in persimmon cultivars were independent of genome-wide structure, and that there was a variety of differentiation paths.

**Fig. 4.**
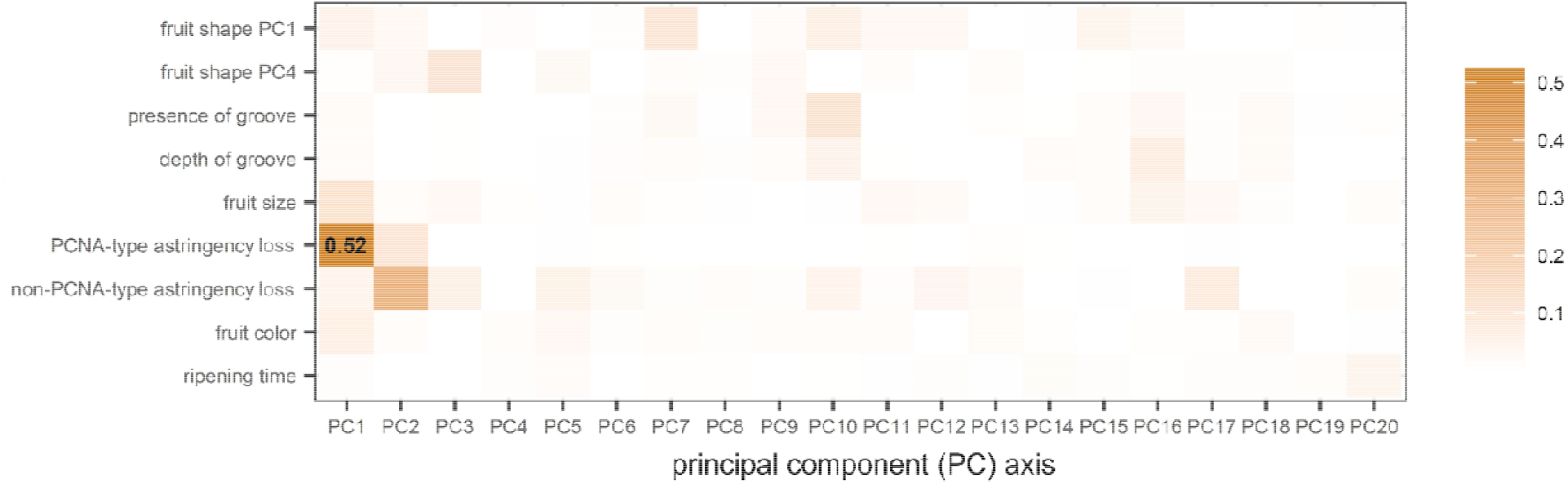
Correlation between population structures and each fruit trait. Pearson’s product-moment correlation (*r*^2^ values) between persimmon fruit traits and the principal components (genome PC1–PC20) of genome-wide SNPs. The PCNA-type astringency-loss trait was substantially correlated with the first principal component, which is consistent with empirical knowledge about persimmon breeding (Sato and Yamada 2016).

### Genomic regions associated with persimmon-specific fruit traits

We conducted GWASpoly to detect associations between genomic regions and the nine traits examined using two types of models (Fig. 2). Our results exhibited significant peaks in fruit shapes and astringency-loss traits in both the dominant and additive models (Fig. 5).

**Fig. 5.**
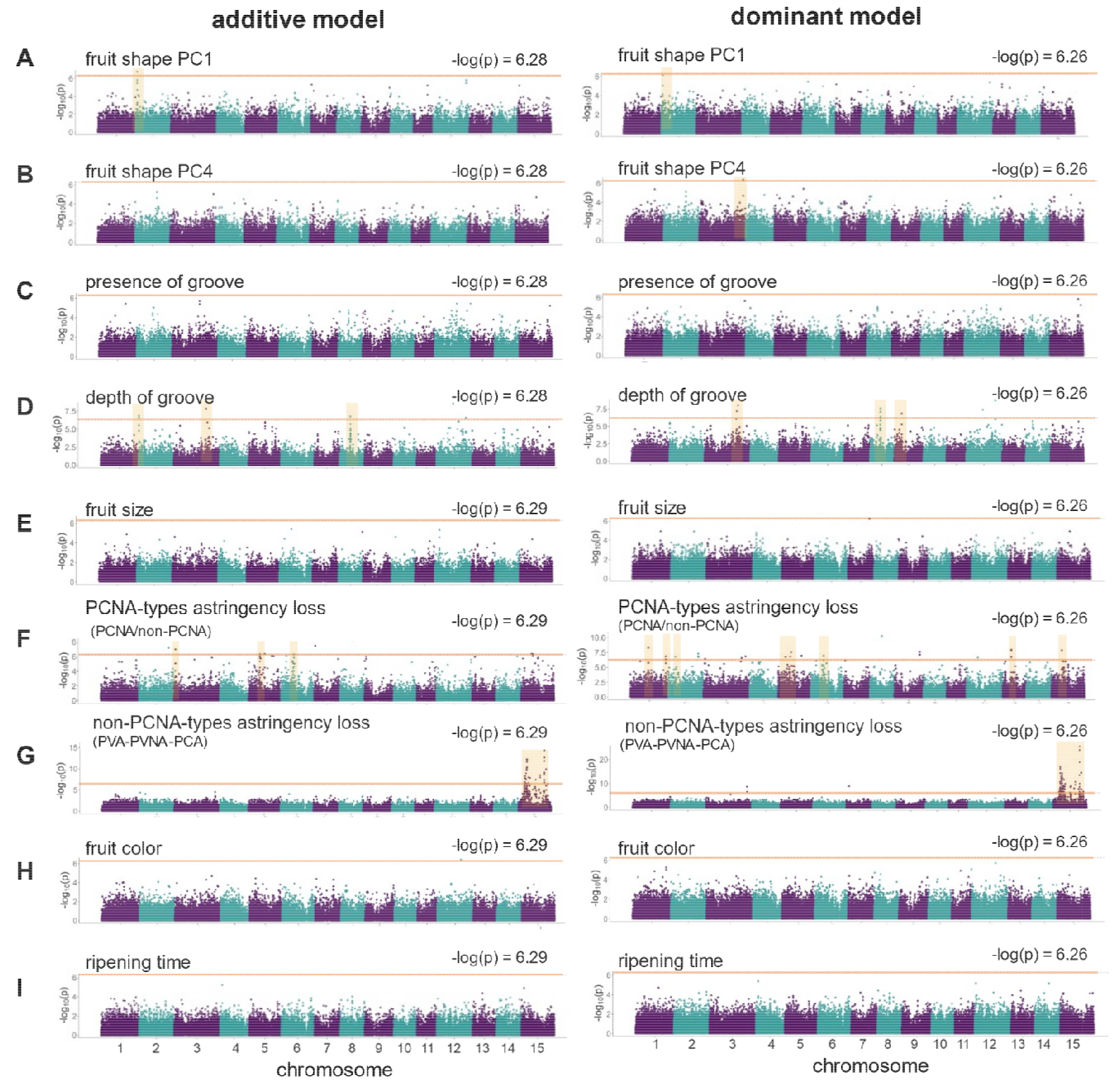
GWAS for representative fruit traits in hexaploid persimmon cultivars. GWAS with additive (left) and dominant (right) models for nine fruit traits in hexaploid persimmon cultivars: fruit-shape PC1 and PC4 values (A and B), presence of groove (C), depth of groove (D), fruit size (E), PCNA-type astringency loss (F), non-PCNA-type astringency loss (G), fruit color (H), and ripening time (I). Potential peak regions are highlighted in pale orange. The thick orange lines indicate −log(*P*-value)>6.26–6.29, which represents the Bonferroni-corrected threshold of 0.1.

For the fruit-shape-related traits, we detected significant associations with fruit shape PC1, fruit shape PC4, and depth of groove, although they are independent of each other. This suggested that persimmon fruit-shape variation is complex and determined by multiple genetic factors. For fruit shape PC1, the highest peak appeared on chromosome 2 in both the additive and 1-dominant models (Fig. 5A). Cultivars with narrow fruit shapes (fruit shape PC1>0.30), except ‘Fudegaki’ (fruit shape PC1=0.40), were heterozygous for the genotype at these loci (Supplementary Fig. S3A). A previous physiological assessment suggested that the narrow (or high fruit shape PC1) fruit shape in ‘Fudegaki’ is due to a unique mechanism: a reduction in the number of locules (Maeda et al. 2018). The fruit shape PC1 value tended to be increased in proportion to allele dosage (Supplementary Fig. S3A), suggesting that the responsible gene (or mutation) can act additively. Linkage disequilibrium was maintained over 1.6 Mbp in the region surrounding the chromosome 2 peak (Supplementary Fig. S4A). Chromosome 12 exhibited a second highest peak associated with fruit shape PC1, although this was slightly less significant in the Bonferroni threshold analysis (Fig. 5A). LD was potentially conserved over 3 Mbp surrounding the peak (Supplementary Fig. S4B). For fruit shape PC4, which is thought to reflect a blossom end shape diversity (Maeda et al. 2018), a peak was observed on chromosome 3 but only in the dominant model (Fig. 5B). For groove depth, significant peaks appeared on chromosomes 3 and 8 in both the dominant and additive models (Fig. 5D), although the peak on chromosome 3 did not overlap the peak associated with fruit shape PC4 (Fig. 5B). Chromosomes 2 and 9 also exhibited significant peaks in the additive and dominant models, respectively (Fig.5D). Although the peak on chromosome 2 for groove depth (Chr2:490,014) was located near a peak for fruit shape PC1 (Chr2: 1,565,911), they were 1 Mbp apart and exhibited no correlation in genotypes (*r*=0.07).

For the two types of astringency-loss traits, the association test for non-PCNA-type astringency loss exhibited a very clear peak on chromosome 15 in both the additive and dominant models (Fig. 5F). A high portion of individuals with astringency loss (or PVNA/PVA traits) tended to be heterozygous for the peak (Supplementary Fig. S5), suggesting that astringency loss is a dominant mutation that establishes a new function: the production of ethanol in seeds to insolubilize high levels of proanthocyanidins in fruit flesh. This is also supported because close wild relatives of hexaploid *D. kaki, D. lotus*, and *D. oleifera* cannot insolubilize proanthocyanidins in fruit. The association test for PCNA-type astringency loss exhibited multiple small peaks scattered throughout the genomes in both the additive and dominant models. Of them, the most promising peak would be on chromosome 5, since an *ASTRINGENCY* locus-linked genomic marker overlapped on the peak in chromosome 5 (*ca*. 29Mb, referring Akagi et al 2010). The lack of a single, clear peak on chromosome 5 might be due to three possible reasons, (i) the clusterized population structure in the PCNA cultivars, as shown in Fig. 3, (ii) high diversities of functional *ASTRINGENCY* alleles (Akagi et al. 2012), and (iii) potential multiple haplotypes of the mutated *ASTRINGENCY* alleles (or *ast*) among the cultivars (Akagi et al. 2012; Kono et al. 2016).

### Genomic regions associated with gene expression involved in fruit-shape diversity

Previous co-expression network analysis studies in persimmon cultivars have highlighted the importance of expression variations of *KNAT1, STK*, and *CKX5* in fruit-shape determination (Maeda et al. 2019). We conducted eGWAS to detect loci potentially associated with the expression levels of these three genes using total 65 cultivars. Note that this expression (transcriptomic) data was sampled over multiple years, and environmental effects might largely affect the results, so here we provided them as supplementary data. In the dominant model, eGWAS showed no peaks for any of the three genes (Supplementary Fig. S6). In the additive model, *STK* and *CKX5* exhibited many noise peaks, potentially because of the small sample size (*N*=65) and substantial environmental effects. On the other hand, *KNAT1* showed a potential peak on chromosome 3 (Supplementary Fig. S6A). Since *KNAT1* (Dlo_pri0307F.1_g01640.1) is located on chromosome 5, it was suggested that the *KNAT1* expression variation in fruit development is regulated not by *cis*-variations but by some mutations in a *trans*-factor, such as a transcription factor, potentially on the chromosome 3 peak, which is supported by the previous studies (Maeda et al. 2019). This peak overlapped with that for fruit groove depth (Fig. 5D), and their genotypes within the peaks were consistent.

### Selective sweep in persimmon cultivar differentiation

SFS-based detection of selective sweep identified statistically significant multiple peaks on chromosomes 3, 4, and 9 (*p*<0.01, Fig. 6). None of them overlapped the peak regions detected by GWAS. This suggests that the loci responsible for hexaploid persimmon-specific fruit traits were not established via strong selections (or bottlenecks) on recent mutations but via natural population transitions. It is worth noting that perennial or tree crops often maintain favored alleles heterozygously during clonal propagation (Numaguchi et al. 2020). Extended haplotype homozygosity (EHH)-based methods (Sabeti et al. 2002, 2007) would better fit this situation than SFS-based methods. However, it would be difficult to properly define haploblocks (or LD decay), as indicated in the method section, in autohexaploid persimmon using only binary SNPs. Thus, further analyses would be required to detect selective sweeps involving persimmon-specific fruit traits.

**Fig. 6.**
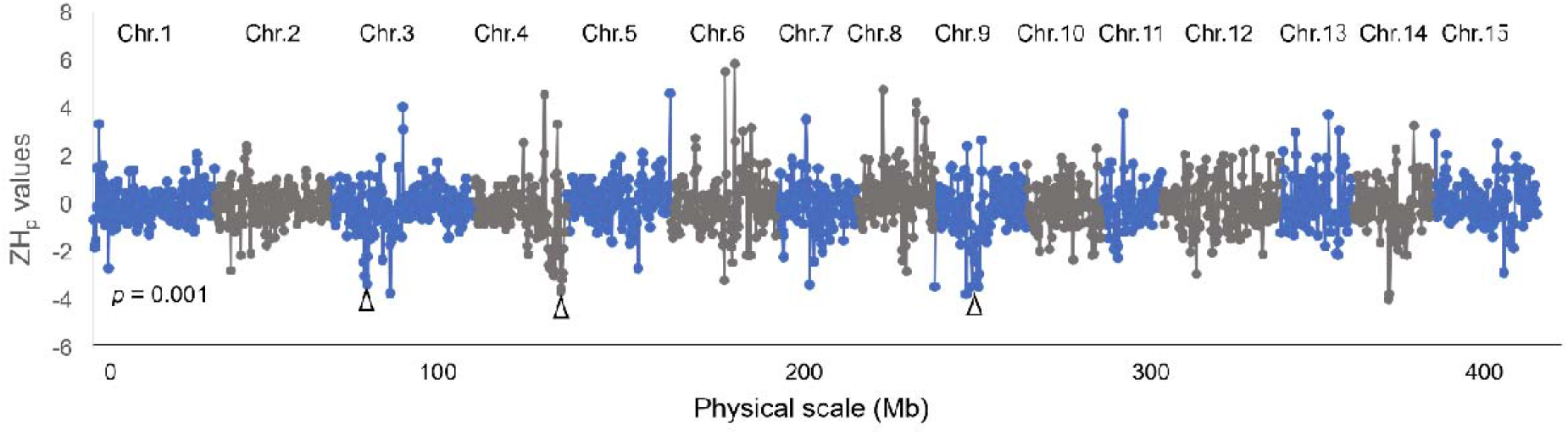
Detection of potential selective sweeps based on a site-frequency spectrum index. Transitions of standardized log-transformed values of the pooled heterozygosity (*ZH*_*p*_; Rubin et al. 2010) in 400-kb bins with 200-kb steps are given. The dotted line indicates the threshold, with *p*<0.001. The peaks exhibiting a gradual reduction in Z*H*_*p*_, which was lower than the threshold, are marked with white arrowheads as potential sweep regions.

## Discussion

### Fruit-shape diversification after hexaploidization

Compared with hexaploid persimmon cultivars, its close diploid wild (or semi-cultivated) relatives, including *D. oleifera* and *D. lotus*, exhibit much less morphological diversity in fruits (Utsunomiya et al. 1998; Yang et al. 2015; Maeda et al. 2018, 2019). This leads to two hypotheses about how fruit-shape diversification was triggered: (i) in pathways arising from domestication (or cultivation) or (ii) via hexaploidization. Our results indicated that the population structure of hexaploid persimmon cultivars is little clustered, except PCNA group (Fig. 3), and there were no principal component axes significantly correlated with the fruit-shape indexes except for fruit astringency traits (Fig. 4). These findings suggested that most of the persimmon cultivars were randomly evolved rather than forming subpopulations (or subclusters), where fruit shapes were also evolved independently; this contrasts with the morphological evolution of grapevine leaves, which is correlated with the transition of population structures (Chitwood et al. 2014). On the other hand, there have been no direct clues or molecular mechanism to explain the hypothesis (ii). Our results so far suggested that the fruit-shape variations of hexaploid persimmon are likely to be controlled by multiple independent genes. Our GWAS analysis detected a locus related to fruit-shape PC1 on chromosome 2 (Fig. 5A), whereas this locus explained only some of the varieties with oblong fruit shapes. In other words, this variety would have lineage-specifically evolved newly arisen gene function (or mutation) with the recent standing variations. Thus, at least for this case, hexaploidization might not be a direct effect but satisfy potential requirements for the current fruit-shape diversity, while local selections of different fruit shapes might contribute more to the current diversity. The loci associated with *KNAT1* expression, which may be responsible for fruit-shape PC1 in a wide variety of persimmon cultivars (Maeda et al. 2019), were detected on chromosome 3 (Fig. 6). Furthermore, other fruit-shape traits, such as the depth of lateral grooves, were associated with chromosome 8, which was independent of fruit-shape PC1 (Fig. 5A-D). Together, these results imply that various genetic factors have accumulated multidimensionally to determine fruit shape among a wide variety of hexaploid persimmon cultivars.

Several examples of the facilitation of phenotypic variations via polyploidization have been reported; for example, studies of *Fragaria* have indicated that polyploids show greater environmental adaptation than diploids (Wei et al. 2018). The results in *Fragaria* support the ‘jack-and-master’ strategy, where non-directional phenotypic changes appear via polyploidization with the potential to be used in adaptation to new environments. *Allium* has also shown higher speciation rates with more trait diversification via intraspecific polyploidy than in diploid or low-ploid populations (Han et al. 2020). Hexaploid persimmon cultivars could be another example of this phenomenon. Considering the phenotypic variations in fruit shape, which are not expressed in close diploids and not correlated with paths of variety differentiation in hexaploids, the ‘jack-and-master’ hypothesis may also be applicable to persimmon, although further experimental validation is necessary. However, the significance of fruit-shape variations in adaptation may be difficult to define, in contrast with other fruit traits, such as astringency traits (or accumulation of second metabolites) or fruit size. Although the SFS-based selective sweep did not overlap with the fruit-shape-associated loci in this study, artificial selections (or bottlenecks) via domestication (or local selections of specific varieties) may be partially involved in development of the current fruit-shape variations, in addition to the effects of hexaploidization.

### Identification of candidate genes determining fruit shape in persimmon

As described, the fruit-shape variations in hexaploid persimmon are thought to be regulated by multiple genetic factors and to have evolved independently (Maeda et al. 2019). Previous report suggested that *KNAT1* is one of main regulators for the fruit shape diversity although the fruit shapes of ‘Saijo’ and its related varieties with high fruit-shape PC1 values are not controlled by *KNAT1* (Maeda et al. 2019). Our GWAS identified the highest peaks on chromosome 2 in both the dominant and additive models (Fig. 5A), and only ‘Saijo’ and its close varieties with high fruit-shape PC1 values showed heterozygous genotypes (Supplementary Fig. S3). Additionally, our eGWAS results identified no peaks on chromosome 2 for *KNAT1* expression (Fig. 6). These results support the abovementioned possibility of fruit-shape diversification via hexaploidization and suggest that candidate fruit shape regulators specific to ‘Saijo’ relatives might be able to be detected. Twenty-nine genes were located within 1.6 Mbp near the peak of chromosome 2, including the gene encoding auxin-inducible in root cultures-like protein, which is involved in auxin production in roots (Gibson and Todd 2015), and that encoding the HAUS augmin-like complex subunit, which is involved in the formation of spindle threads during cell division (Hotta et al. 2012; Lee et al. 2017). Since the fruit-shape variation in persimmon cultivars depends on diversity in cell division (or proliferation) in the mesocarp (Maeda et al. 2018), these genes may be good candidates as determinants of the specificity of fruit shape in the ‘Saijo’ variety. Chromosome 12 also exhibited an association peak for fruit-shape PC1, independent of the peak on chromosome 2 (Fig 5A). On chromosome 12, 244 genes, including *CYCD*, which is involved in the cell cycle, were localized to regions surrounding the peaks with significant LD values. On chromosome 3, the peaks associated with groove depth and *KNAT1* expression overlapped. This suggested that a gene regulating *KNAT1* that also causes cell division defects in the locule marginal regions, which results in peripheral fruit grooves, is a candidate. Our results successfully identified the complicated genetic bases for fruit-shape variations in hexaploid persimmon. Additional fine-expression profiling covering the entire process of fruit development in specific genotypes identified in this study will narrow down the candidate genes. It is also worth noting that this study referred to the genome information of a close diploid relative, *D. lotus*, which shows no substantial fruit-shape variations and has no genes/mutations corresponding the fruit-shape diversity observed in hexaploid *D. kaki* cultivars. Hence, it would be much preferable to use the genome information of the hexaploid *D. kaki* itself when discussing the possibility of fruit shape diversification through hexaploidization.

## Acknowledgments

We thank Candace Webb, Ph.D., from Edanz (https://jp.edanz.com/ac) for editing a draft of this manuscript. This work was supported by PRESTO from Japan Science and Technology Agency (JST) [JPMJPR20Q1] and Grant-in-Aid for Scientific Research (B) [22H02339] and for Transformative Research Areas (A) [22H05172 and 22H05173] from JSPS to T.A.

## Author contribution statement

T.A. conceived the study. A.H., K.M., and T.A. designed experiments. A.H., K.M., and K.S. conducted the experiments. A.H., K.M., and T.A. analyzed the data. N.O., R.M., R.T., Y.K., K.U., and T.A. contributed to plant resources and facilities. A.H., and T.A. drafted the manuscript. All authors approved the manuscript.

## Data archiving

The sequence reads are available from the DDBJ Sequence Read Archive (DRA) under the accession number DRA015334.

