## Supplementary Figures S1-S6 for "Genetic basis of lineage-specific evolution of fruit traits in hexaploid persimmon"

**Supplementary Fig. S1**

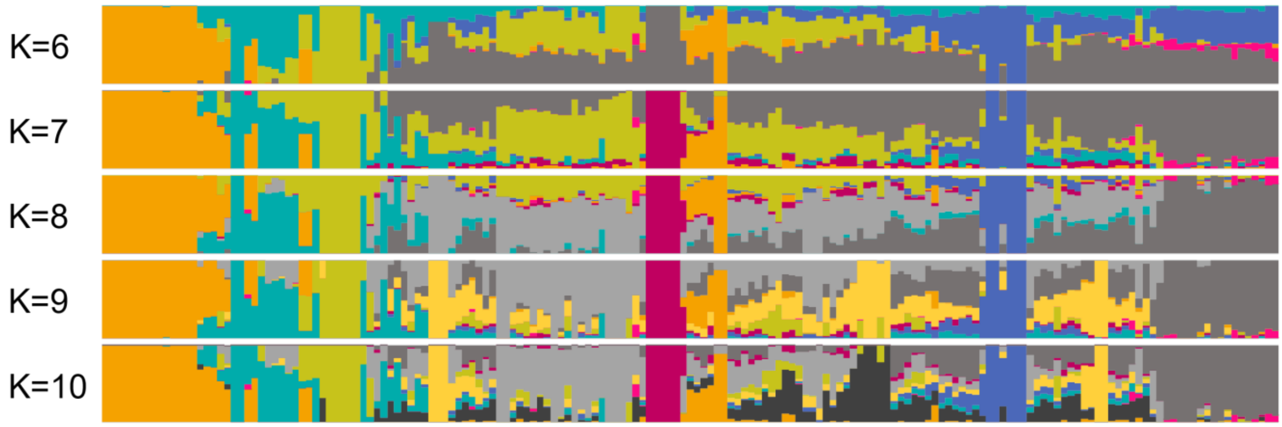


**Supplementary Fig. S1 ADMIXTURE in hexaploid persimmon cultivars with K=6-10.** Proportion of ancestry from K=6–10 inferred using ADMIXTURE in 173 hexaploid persimmon cultivars. The orange circles indicated PCNA cultivars. PCNA-type astringency-loss varieties mostly formed a separate cluster. The cultivars used in ADMIXTURE were listed in Supplementary Table S3.

**Supplementary Fig. S2**

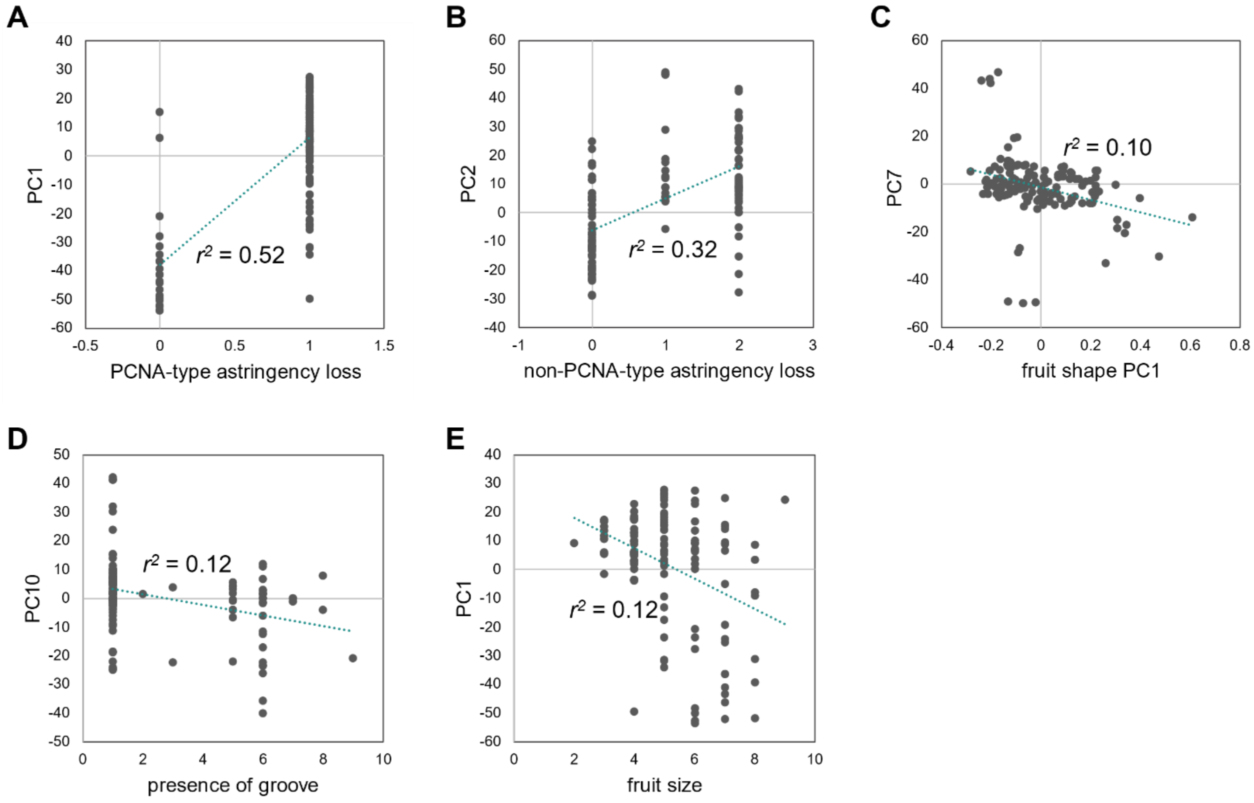


**Supplementary Fig. S2 Correlations between the genome structure PCs and the fruit phenotypic values.**

From the comprehensive correlation between the genome structure PC1-20 and the nine fruit traits (Figure 4), we visualized 5 correlations with significant correlation coefficients ($|r| > 0.3$). (A) genome PC1 and PCNA-type astringency loss, (B) genome PC2 and non-PCNA-type astringency loss, (C) genome PC7 and fruit shape PC1, (D) genome PC10 and presence of groove, (E) genome PC1 and fruit size. The slight correlation in genome PC7 and fruit shape PC1 (C) might be due to the outliers.

**Supplementary Fig. S3**

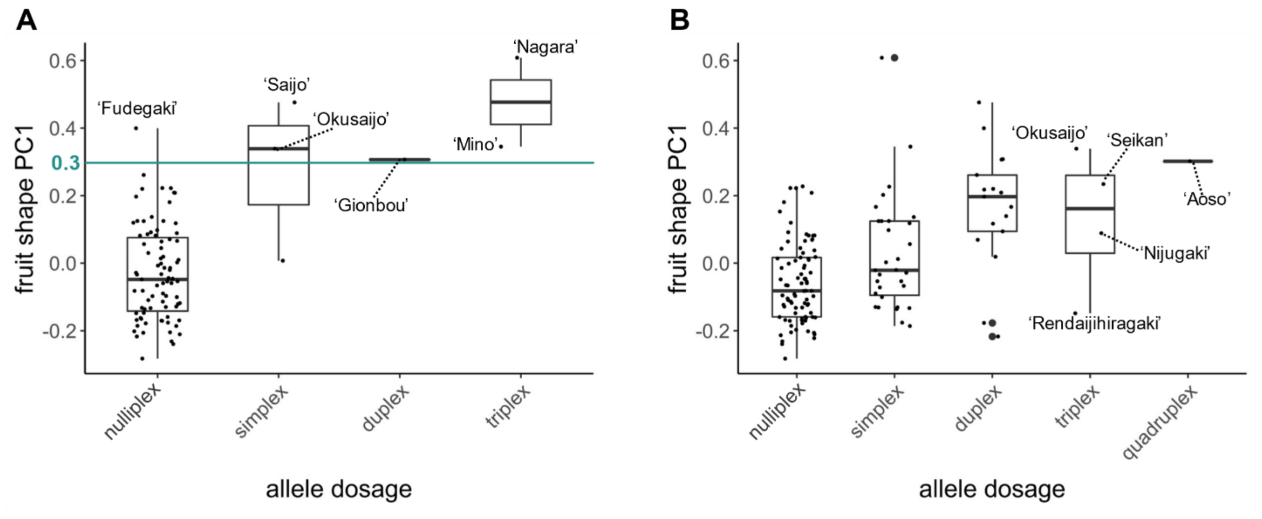


**Supplementary Fig. S3 Fruit shape PC1 distribution in the quantitative genotypes of the association peaks in chromosomes 2 and 12.**

Fruit shape PC1 distributions in nulliplex to quadruplex allele states at the peaks of chromosome 2 (A) and chromosome 12 (B). The individuals with heterozygous genotypes (or simplex-quadruplex) at the peaks of chromosomes 2 and 12 were independent (r = 0.27), suggesting that they are distinct genetic factors affecting fruit shape PC1.

**Supplementary Fig. S4**

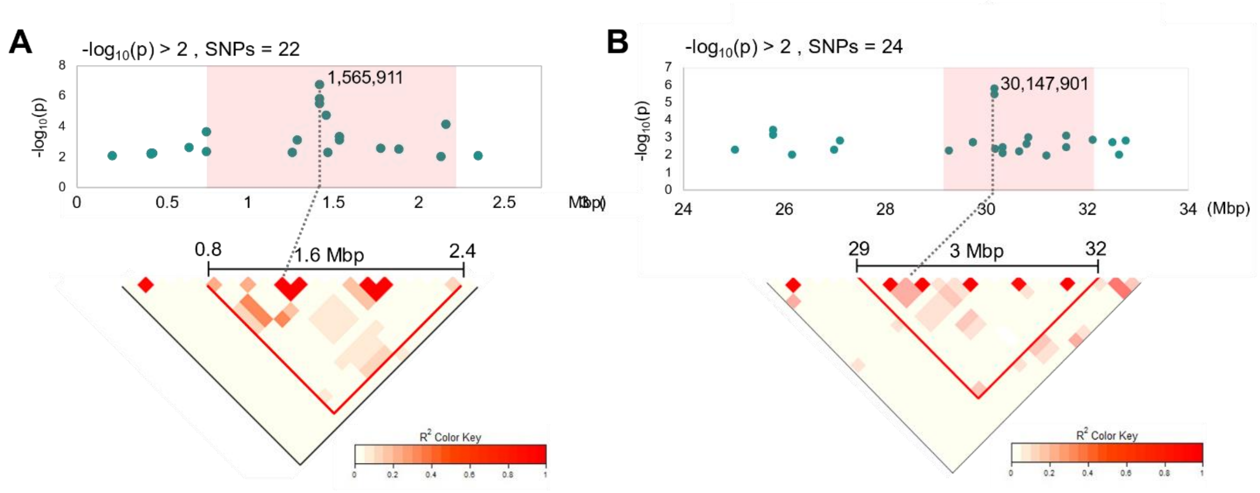


**Supplementary Fig. S4 LD conservation surrounding the GWAS peaks at chromosomes 2 and 12, for association with fruit shape PC1.**

From the highest summits (Chr2: 1,565,911, and Chr12: 30,147,901), approximately 1.6Mb and 3Mb LDs were maintained, respectively. The discontinuous LD decays would not reflect complex standing genetic variations in persimmon but are mainly due to technical issues in autohexaploid (or autopolyploid) genotypes, which can define multiple heterozygous states (simplex-pentaplex), in contrast to diploids.

**Supplementary Fig. S5**

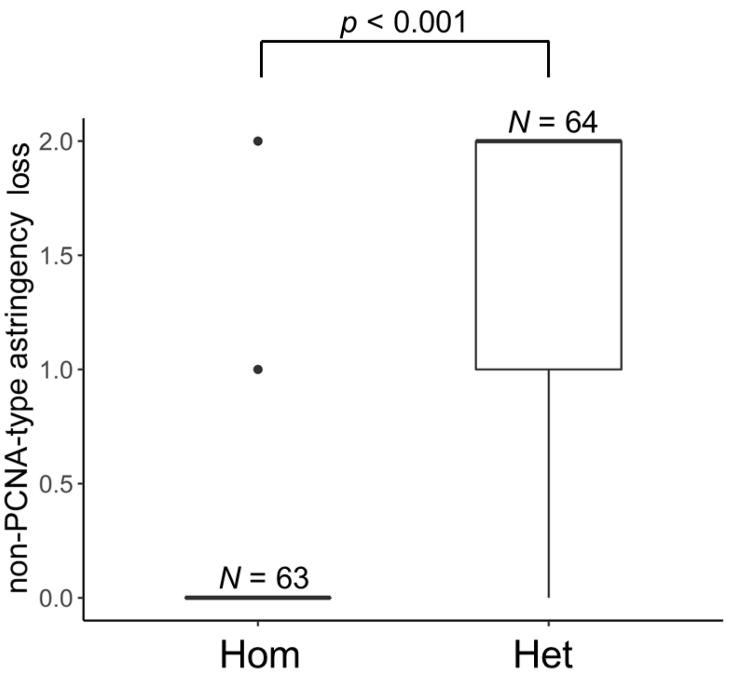


**Supplementary Fig. S5 Distribution of the phenotypic values for non-PCNA-type astringency-loss in the quantitative genotypes of the association peak in chromosome 15**

Most of the astringent types (or pollination-constant astringent: PCA) showed homozygous (or nulliplex in hexasomic genotype) for the genotype at the peak of chromosome 15. Considering the close wild relatives of *D. kaki*, such as *D. lotus* or *D. oleifera*, exhibit constant astringency in the fruit, this result indicates that non-PCNA type astringent loss is a dominant new function established in a lineage-specific manner in hexaploid *D. kaki*.

**Supplementary Fig. S6**

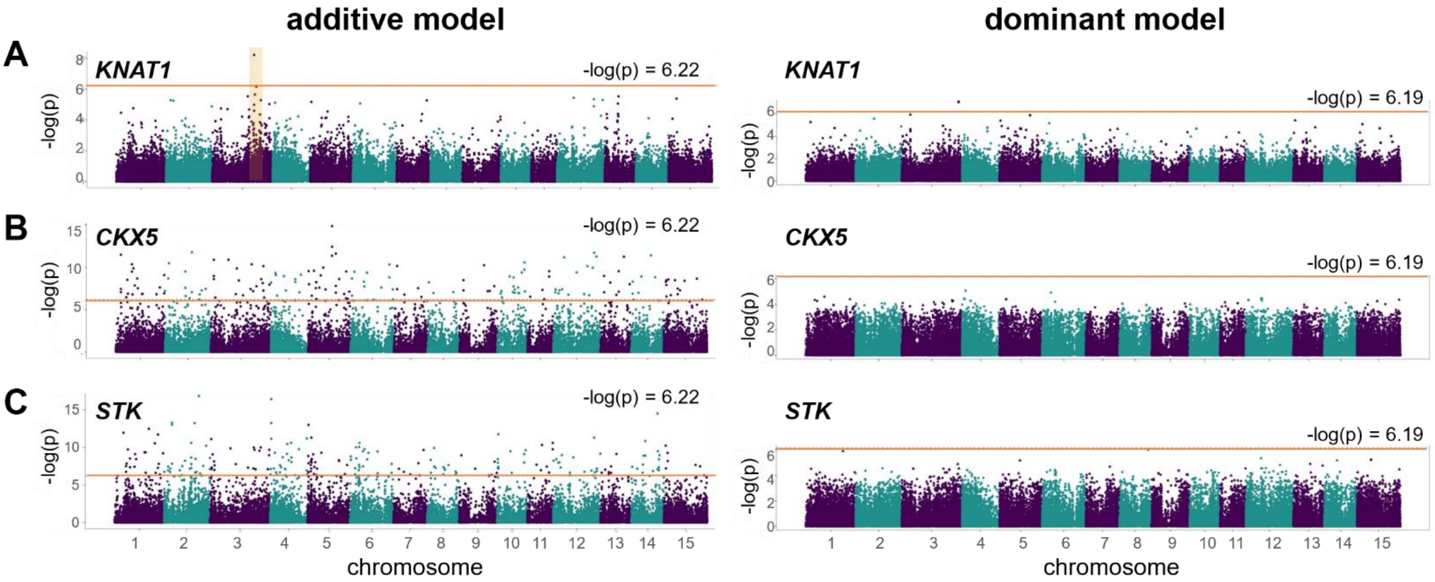


**Supplementary Fig. S6 eGWAS for the three genes potentially important for persimmon fruit shape diversity.**

eGWAS with additive model (left) and dominant model (right), for expression levels of the three genes, associated with persimmon fruit shape, *KNAT1* **(A)** *CKX5* **(B)**, and *STK* **(C)** (Maeda et al. 2019). Potential peak regions are highlighted in pale orange. The thick orange lines indicate −log(*P*-value) > 6.19-6.22, representing the Bonferroni-corrected threshold of 0.1.
